# Poor neutralizing antibody responses against SARS-CoV-2 Omicron BQ.1.1 and XBB in Norway in October 2022

**DOI:** 10.1101/2023.01.05.522845

**Authors:** Elisabeth Lea Vikse, Even Fossum, Magnhild Sekse Erdal, Olav Hungnes, Karoline Bragstad

**Author notes:** These authors contributed equally to this work and share first authorship (Elisabeth Lea Vikse, Even Fossum).

## Abstract

New sub-lineages of the SARS-CoV-2 omicron variants with enhanced ability to evade existing antibody responses continue to evolve. A better understanding how susceptible emerging virus variants are to immunity induced by vaccination or infection could help predict which strains will become dominant going forward. Here we evaluate neutralizing antibodies against several clinical isolates of omicron variants including BQ.1.1 and XBB in sera from 3x mRNA vaccinated individuals and individuals with breakthrough infections with early (BA.1 or 2) or late (BA.5) omicron variants. In addition, we evaluate neutralizing antibodies in serum samples harvested from 32 individuals from the middle of October 2022, to provide a more recent estimate of immunity. As expected, serum samples harvested after breakthrough infections were more efficient at neutralizing all the omicron variants, compared to sera from non-infected individuals. While neutralization remained high against variants such as BA.2.75.2, BR.1 and BF.7, there was a marked reduction in neutralizing titers against BQ.1.1 and XBB. Similarly, most serum samples harvested in October 2022 had very low neutralizing antibodies against BQ.1.1 and XBB, suggesting that these variants and their descendants will dominate infection waves in Norway this winter season.

## Introduction

New Omicron subvariants have demonstrated increased ability to evade immune responses induced by vaccination and/or infection[1–3], and resistance to existing monoclonal antibody treatments such as Evusheld (tiksagevimab and cilgavimab) and bebtelovimab [4]. Immune evasion has been linked to several mutations occurring in the receptor binding domain (RBD) of the spike surface antigen, including aa positions R346, K444, L452, N460 and F486 [2, 5, 6]. Recently, sub-lineages of both BA.2.75 and BA.5 have independently acquired substitutions in these aa positions, suggesting converging evolution and growth advantages relative to non-mutated variants [7, 8]. While early variants of BA.2.75 and BA.5 contained one or two mutations in this group of aa positions, the more recently described pango lineage BQ.1.1 (BA.5 derived) have acquired substitutions in all 5 aa positions. In addition, recombination of two BA.2 derived lineages (BJ.1 and BM.1.1.1) has resulted in the formation of the XBB variant which has proven highly immune evasive [6]. The XBB variant contains substitutions in three of the RBD positions (R346, N460 and F486) in addition to a deletion in the N-terminal domain of spike S1 (Y144).

Prior to the emergence of the omicron variant in November 2021 [9], infections with SARS-CoV-2 remained relatively low in Norway due to non-pharmaceutical restrictions and a high vaccination coverage. In August 2021, a total of ~150.000 infections had been registered, corresponding to about 3 % of the population. However, seroprevalence of IgG directed against nucleoprotein (N), an antigen not included in any of the vaccines used in Norway, was determined to be 11.7 % in August 2021, suggesting that the number of infected was significantly higher [10]. Nevertheless, most Norwegians remained uninfected prior to omicron.

By the end of September 2022, >89% of Norwegians over 18 years had received at least two doses of covid-19 vaccine, and >91 % of those over 65 years had received at least three doses [11]. Combined with the introduction of the Omicron variant in November 2021 [12], and the subsequent BA.1/BA.2 wave in January - March and BA.4/BA.5 wave in summer 2022, the vast majority of Norwegians has acquired immune responses against SARS-CoV-2. It is, however, uncertain how well these responses protect against novel Omicron variants. To remedy this situation, we have evaluated neutralizing antibodies against isolates of recent Omicron variants in sera from recipients of 3 doses of monovalent mRNA vaccine, and from individuals with BA.1, BA.2 or BA.5 breakthrough infections. In addition, we evaluate neutralizing antibodies against BA.5, BQ.1.1 and XBB in serum samples harvested from 32 individuals in October 2022.

## Results

### Immune evasion of new omicron variants

To evaluate neutralizing antibodies against variants of interest, isolates of Omicron variants BA.2, BA.5, BA.2.75, BA.2.75.2, BF.7, BR.1, BQ.1.1 and XBB were grown in VeroE6/TMPRSS2 or Vero/hSLAM cells (Supplementary Table I). Throughout the pandemic, SARS-CoV-2 specimens have been genetically characterized by whole genome sequencing as part of the Norwegian covid-19 surveillance effort. Specimens containing variants with combinations of mutations reported to induce immune escape have been selected for virus isolation (Figure 1A and Supplementary Table I). Virus isolates were passaged twice, and the second passage used for the subsequent neutralization assays. The passaged virus was sequenced to ensure the genotype of the variant. While we observed a limited number of nucleotide substitutions in the passaged viruses, none led to aa changes in the Spike protein.

**Figure 1:**
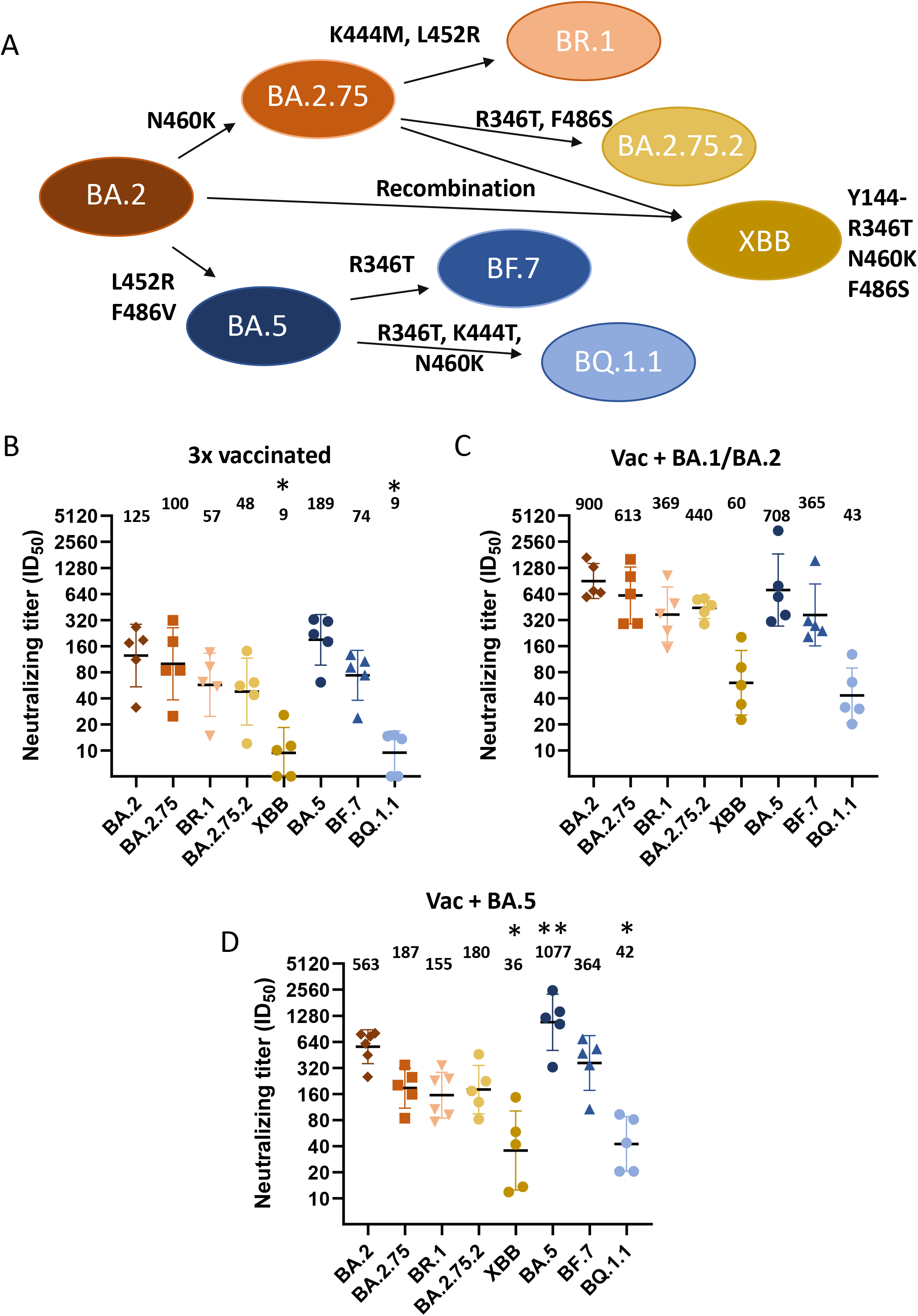
Omicron variants evade neutralizing antibodies induced by vaccination and breakthrough infection. A) Overview of Omicron variants evaluated by neutralization assay, including the most relevant aa substitutions in the RBD of spike. B-D) Neutralizations assays using sera from three times vaccinated individuals without breakthrough infection (B), or from vaccinated individuals with breakthrough infection with BA.1 or BA.2 (C) or BA.5 (D). Geometric mean is indicated above each dataset. Data presented as geometric mean ± geometric standard deviation. N = 5 donors for each dataset. One-way ANOVA compared to BA.2 with Dunnett’s multiple comparison test, * = p < 0.05, ** = p < 0.01.

Sera from three times mRNA vaccinated individuals and from two- or three-times vaccinated individuals with subsequent breakthrough infections with BA1, BA.2 or BA.5 were analyzed for the ability to neutralize the viral isolates. Serum was harvested 3 – 4 weeks after vaccination or breakthrough infection. While sera from three times vaccinated individuals neutralized BA.2 and BA.5 isolates well, there were a slight reduction against BR.1, BA.2.75.2 and BF.7. Neutralizing titers were, however, significantly reduced against BQ.1.1 and XBB, (Figure 1B).

Sera from people with breakthrough infections with BA.1, BA.2 or BA.5 generally had higher neutralizing titers against all tested variants, compared to vaccinated-only individuals (Figure 1C-D). As expected, people with BA.5 breakthrough infection had significantly higher neutralizing titers against BA.5 virus, compared to BA.2 virus (Figure 1D). In contrast, people with BA.1 or BA.2 breakthrough infections had similar neutralizing titers against both BA.2 and BA.5, and strong neutralizing titers against more recent variants such as BA.2.75, BR.1, BA.2.75.2 and BF.7 (Figure 1C). There was, however, a marked reduction in neutralizing antibodies against the BQ.1.1 and XBB in people with breakthrough infection (Figure 1C-D).

### Protection against BQ.1.1 and XBB in October 2022

Serum samples taken shortly after vaccination or breakthrough infection may not necessarily provide a good picture of the current levels of neutralizing antibodies. To evaluate immunity towards the more immune evasive Omicron variants prior to the winter season, 32 serum samples were harvested from healthy individuals on 13-14 October 2022, and evaluated for neutralizing titers against BA.5, BQ.1.1 and XBB. The average age of the donors were 42.6 years, and the donors consisted of 78% females. While neutralizing titers against BA.5 remained elevated (≥64) in 20 of 32 tested individuals, there was a significant reduction in neutralizing antibodies against BQ.1.1 and especially XBB (Figure 2A). For BQ.1.1, 9 of 32 had neutralizing titers >64, and only 4 of 32 had titers >64 against XBB.

**Figure 2:**
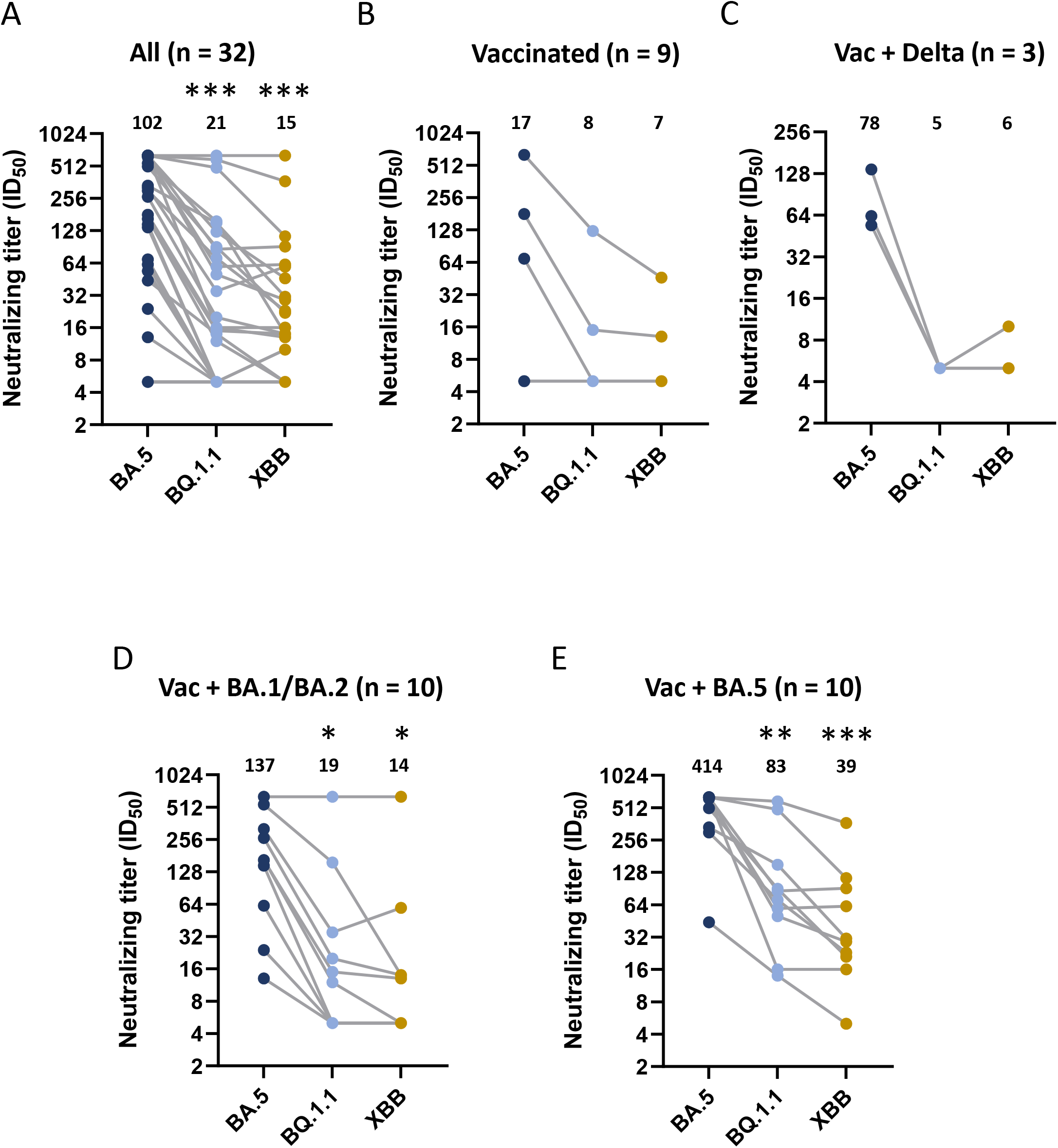
Neutralizing antibodies towards BQ.1.1 and XBB in sera collected October 2022. A) Sera from 32 donors were evaluated in neutralization assays against Omicron variants BA.5, BQ.1.1 and XBB. B-D) Sera from (A) were divided into groups of vaccinated without infection (n = 9) (B), vaccinated with delta breakthrough infection (Vac + Delta, n = 3) (C), vaccinated with BA.1 or BA.2 breakthrough infection (Vac + BA.1/BA.2, n = 10) (D) or vaccinated with BA.5 breakthrough infection (Vac + BA.5 n = 10) (E), and evaluated for neutralizing antibodies against BA.5, BQ.1.1 or XBB. Geometric mean is indicated above each dataset. Data presented as geometric mean ± geometric standard deviation. (A) n = 32, and (B-E) n = 3-10 donors for each dataset. Matched One-way ANOVA with Tukey’s multiple comparison test, * = p < 0.05, ** = p < 0.01, *** p < 0.001.

Next, we divided the donors into groups based on vaccination and infection history (Supplementary Table II). It should be noted that donors may have undergone mild or subclinical infections which have not been accounted for. In general, presumed uninfected donors had lower titers of neutralizing antibodies against all tested variants (Figure 2B). One uninfected donor had recently received a 4^th^ vaccine dose and had the highest titers against all variants in this group. Donors with previous breakthrough infections with BA.1 or BA.2 and especially BA.5, had high neutralizing titers towards BA.5. Interestingly, the tree donors with only Delta breakthrough also had reasonably high titers against BA.5 (Figure 2C). There were no non-vaccinated individuals among the donors.

There was a significant reduction in neutralizing antibodies against BQ.1.1 and XBB in donors with BA.1, BA.2 or BA.5 breakthrough infections (Figure 2D-E). However, 6/10 donor with BA.5 breakthrough infection maintained neutralizing titers >64 (Figure 2E), which may provide some protection from infection [13]. People with BA.5 breakthrough had higher neutralizing titers against BQ.1.1 and XBB, than people with BA.1 or BA.2 breakthrough, but this is likely also due to the shorter time since infection.

## Discussion

Here we present data on neutralizing antibodies against various SARS-CoV-2 Omicron variants in serum samples from Norwegian donors in October 2022. There was a significant reduction in neutralizing antibodies against the more immune evasive BQ.1.1 and XBB variants. However, neutralization was stronger in people who had undergone recent BA.5 breakthrough infections.

Our observations corroborate the immune evasive nature of the BQ.1.1 and XBB variant as observed by others in pseudovirus-neutralization assays [7] and live virus assays [6, 14]. When evaluating sera harvested 3-4 weeks after the third vaccination or after breakthrough infection, neutralizing antibody levels were similar against BQ.1.1 and XBB. Donors with breakthrough infection of BA.1 or BA.2 even had higher neutralizing titers against XBB than BQ.1.1, which is in contrast to what others have observed [6, 14]. When evaluating neutralizing titers from sera harvested in October 2022, donors with breakthrough infections with BA.5 had higher neutralizing titers against both BQ.1.1 and XBB compared to donors with BA.1 or BA.2 breakthrough. However, this likely reflects the more recent BA.5 wave of infections, and not that BA.5 infection induces neutralizing antibodies with broader specificity. Indeed, sera harvested 3-4 weeks after BA.1 or BA.2 infection neutralized BA.2.75 derived variants (BR.1 and BA.2.75.2) and BA.5 derived variant (BF.7) equally well, while sera from BA.5 breakthrough neutralized BF.7 better than BR.1 and BA.2.75.2 (Figure 1).

Although neutralizing antibodies have been established as a correlate of protection against SARS-CoV-2 [13, 15, 16], it is less clear what constitutes protective levels of neutralizing antibodies. Gilbert and colleagues observed a 91% risk-reduction of covid-19 (symptomatic infection) with neutralizing titer of 100 in a pseudotype assay during a 100-days follow-up period [15]. Similarly, Dimeglio and colleagues observed 94 % reduced risk of infection (symptomatic and asymptomatic) with a neutralizing titer of 64-128 using live virus, over a follow-up period of 275 days [13]. In our experiments, 9/32 donors had neutralizing antibody titers over 64 against BQ.1.1, including 6/10 donors with BA.5 breakthrough infections. While protective efficacy cannot be extrapolated onto our data, it seems likely that recent BA.5 breakthrough infections confer some protection against BQ.1.1. In contrast, only 4/32 donors had neutralizing titers above 64 against the XBB variant.

## Conclusion

We see a marked reduction in neutralizing antibodies against the immune evasive Omicron variants BQ.1.1 and XBB in serum samples from October 2022, and as of December 2022, BQ.1, BQ.1.1 and descendants are the dominating lineages in Norway [11]. Nevertheless, people with BA.5 breakthrough infections are likely to retain some protection against especially BQ.1.1. The fact that immunity against BA.1/BA.2 has waned over several months, combined with lower neutralizing titers against XBB (BA.2 recombinant) observed here, suggests that this variant and its descendants may also give rise to new waves of infections this winter in Europe.

## Material and Methods

### Sera from donors

Sera was collected from employees at the Norwegian Institute of Public health. All samples were collected with written consent. For Figure 1, sera were harvested 3-4 weeks after vaccination or breakthrough infection with a confirmed BA.1, BA.2 or BA.5 infection. For sera harvested in October 2022, donors were defined as BA.1 or BA.2 breakthrough if they had been infected in January-March, and BA.5 infected if they had been infected during May-August. All previously uninfected individuals were fully vaccinated (3 doses of mRNA vaccines), and one donor had received a 4^th^ dose with a bivalent mRNA vaccine. It should be noted that donors may have undergone mild or subclinical infections which have not been accounted for in the groups.

### Virus isolation and titration

In a biosafety level 3 (BSL3) facility, viral specimens collected from SARS-CoV-2 infected patients in Norway and selected based on the viral genome sequence were incubated on Vero E6/hSLAM (ECACC #04091501) or Vero E6/TMPRSS2 (NIBSC #100978) cells for 1 hour at 37°C. After incubation the inoculate was removed and replaced with fresh culture medium. The cells were incubated for 3-4 days at 37 °C and a small volume of the supernatant was passaged onto fresh cells. After 3-4 more days the second passage of virus was harvested and frozen in aliquots. The viral stocks were analyzed by qRT-PCR and sequenced by Illumina NGS to confirm the variant sequence and to ensure that no critical mutations had been selected for during the virus cultivation in cells. The harvested virus was titrated on Vero E6 cells to determine the tissue culture infectious dose (TCID50) before the virus neutralization assay was performed.

### Neutralization assay

The neutralization assay and ELISA protocol was adapted and modified from Krammer et al. [17]. Post-exposure sera were collected three to four weeks post vaccination or infection. In addition, we collected sera from 32 individuals in October 2022 from both fully vaccinated non-infected individuals and individuals with previous breakthrough infections of various SARS-CoV-2 variants. The negative serum control used was a pool of sera collected in early 2019. The sera were heat inactivated for 30 minutes at 56 °C and serially diluted twofold in a 96 well deep well plate. In a BSL-3 facility, a viral dose of 100×TCID_50_ was added to each well of diluted sera and to a virus control not containing serum. Cell controls only containing virus diluent were also included. The virus-serum mix incubated at 37°C for 1h and was then added to a 96 well plate of Vero E6 cells with 12000 cells per well seeded the day before. After 96 hours of incubation at 37°C the plates were checked for cytopathic effect using a light microscope. The cells were fixed with 80% acetone, and the plates transported out of the BSL3 facility.

An ELISA detecting the nucleocapsid of SARS-CoV-2 was performed on the fixed cell layer in a BSL2 facility. The ELISA consists of a blocking step using PBS with 1% BSA, primary incubation with SARS-CoV-2 nucleocapsid antibody (Sino biological, 40143-R019) and secondary incubation with Goat anti-rabbit IgG Alkaline Phosphatase Antibody (Sigma, A3687).

Between each incubation step the plates were washed with wash buffer (PBS with 2% Tween 20). Finally, 1 mg/mL of phosphatase substrate dissolved in diethanolamine buffer was added to the plates, and the OD405 was measured for each well after 40 minutes. The virus neutralization 50% titer was calculated from the measured OD values and the results were graphed and statistically analyzed using Graph-Pad PRISM version 9.

### Statistics

Neutralization data is presented as geometric mean with geometric standard deviation. One-way ANOVA with Dunnett’s (Figure 1) or Tukey’s multiple comparison test was used to compare data sets. All statistics were calculated using Graph-Pad PRISM version 9.

## Statements

### Ethical statement

A separate ethical approval was not obtained, as this study was a part of the routine surveillance and investigation of virus properties performed at NIPH as the national reference laboratory for coronaviruses with outbreak potential. NIPH is authorized to conduct these studies under the Norwegian Infection control act and the regulations relating to notifiable diseases. All human sera (from colleagues at the NIPH) were obtained with written consent.

### Data availability

Anonymised data can be shared in accordance with the data sharing policy of NIPH. Virus genome sequences (original specimen) are available in GISAID EpiCoV with accession numbers EPI_ISL_12981999 (BA.5), EPI_ISL_16100571 (BA.2), EPI_ISL_14773262 (BF.7), EPI_ISL_14892153 (BA.2.75), EPI_ISL_15191765 (BR.1), EPI_ISL_15349765 (BQ.1.1), and EPI_ISL_15538637 (XBB).

## Acknowledgements

Vero E6/hSLAM cells were kindly provided by the Dutch National Institute for Public Health and the Environment, Bilthoven, The Netherlands, with the kind approval of its originator, Professor Yusuke Yanagi. We would like to thank all the donors for providing serum samples, and Hang Thi Ngoc Le and Julie Tro Jerud for assistance with blood sampling. We acknowledge the national microbiological laboratories in Norway supporting the surveillance of Covid-19 with clinical samples. We thank Ignacio Garcia Llorente and Rasmus Kopperud Riis for deep sequence analysis of the virus isolates, and also thank our other highly skilled colleagues in the Department of Virology and beyond who contributed to make the study possible, e.g. with virus sequencing under the coordination of Kathrine Stene-Johansen and provision of cell cultures supervised by Inger M. Böckerman.

## Authors’ contributions

ELV and MSE planned and performed experiments and analysed data. EF supervised experiments, analysed data and drafted the first manuscript. OH and KB conceptualised the study and supervised experiments. All authors reviewed the manuscript and approved of the final version.

