## Supplementary tables for "Poor neutralizing antibody responses against SARS-CoV-2 Omicron BQ.1.1 and XBB in Norway in October 2022"

**Supplementary table I:** Viral isolates used for neutralization assay. Virus genome sequences (original specimen) are available in GISAID EpiCoV with accession numbers EPI_ISL_12981999 (BA.5), EPI_ISL_16100571 (BA.2), EPI_ISL_14773262 (BF.7), EPI_ISL_14892153 (BA.2.75), EPI_ISL_15191765 (BR.1), EPI_ISL_15349765 (BQ.1.1), and EPI_ISL_15538637 (XBB).

| Isolate | Sample date | Pangolin variant | Defining mutations in RBD | Full list of mutations in spike, relative to nCoV2019/Wuhan-Hu-1/MN908947 |
|---|---|---|---|---|
| SC2/Norway/00126/2022 | 02.01.2022 | BA.2 | Reference | T19I;L24S;G142D;V213G;G339D;S371F;S373P;S375F;T376A;D405N;R408S;K417N;N440K;S477N;T478K;E484A;Q493R;Q498R;N501Y;Y505H;D614G;H655Y;N679K;P681H;N764K;D796Y;Q954H;N969K;P25-;P26-;A27- |
| SC2/Norway/21334/2022 | 02.05.2022 | BA.5 | L452R, F486V | T19I;L24S;G142D;V213G;G339D;S371F;S373P;S375F;T376A;D405N;R408S;K417N;N440K;L452R;S477N;T478K;E484A;F486V;Q498R;N501Y;Y505H;D614G;H655Y;N679K;P681H;N764K;D796Y;Q954H;N969K;D1139Y;P25-;P26-;A27-;H69-;V70- |
| SC2/Norway/29067/2022 | 31.07.2022 | BF.7 | R346T, L452R, F486V | T19I;L24S;G142D;V213G;G339D;R346T;S371F;S373P;S375F;T376A;D405N;R408S;K417N;N440K;L452R;S477N;T478K;E484A;F486V;Q498R;N501Y;Y505H;D614G;H655Y;N679K;P681H;N764K;D796Y;Q954H;N969K;P25-;P26-;A27-;H69-;V70- |
| SC2/Norway/30134/2022 | 16.08.2022 | BA.2.75 | N460K | T19I;L24S;G142D;K147E;W152R;F157L;I210V;V213G;G257S;G339H;K356T;S371F;S373P;S375F;T376A;D405N;R408S;K417N;N440K;G446S;N460K;S477N;T478K;E484A;Q498R;N501Y;Y505H;D614G;H655Y;N679K;P681H;N764K;D796Y;Q954H;N969K;P25-;P26-;A27- |
| SC2/Norway/32397/2022 | 14.09.2022 | BA.2.75.2 | R346T, N460K, F486S | T19I;L24S;G142D;K147E;W152R;F157L;I210V;V213G;G257S;G339H;R346T;S371F;S373P;S375F;T376A;D405N;R408S;K417N;N440K;G446S;N460K;S477N;T478K;E484A;F486S;Q498R;N501Y;Y505H;D614G;H655Y;N679K;P681H;N764K;D796Y;Q954H;N969K;D1199N;P25-;P26-;A27- |
| SC2/Norway/31280/2022 | 09.09.2022 | BR.1 | K444M, L452R, N460K | T19I;L24S;G142D;K147E;W152R;F157L;I210V;V213G;G257S;G339H;S371F;S373P;S375F;T376A;D405N;R408S;K417N;N440K;K444M;G446S;L452R;N460K;S477N;T478K;E484A;Q498R;N501Y;Y505H;D614G;H655Y;N679K;P681H;N764K;D796Y;Q954H;N969K;P25-;P26-;A27- |
| SC2/Norway/31371/2022 | 15.09.2022 | BQ.1.1 | R346T, K444T, L452R, N460K, F486V | T19I;L24S;G142D;V213G;G339D;R346T;S371F;S373P;S375F;T376A;D405N;R408S;N440K;K444T;L452R;N460K;S477N;T478K;E484A;F486V;Q498R;N501Y;Y505H;D614G;H655Y;N679K;P681H;N764K;D796Y;Q954H;N969K;P25-;P26-;A27-;H69-;V70- |
| SC2/Norway/32121/2022 | 05.10.2022 | XBB | Y144-, R346T, N460K, F486S | T19I;L24S;V83A;G142D;H146Q;Q183E;V213E;G339H;R346T;L368I;S371F;S373P;S375F;T376A;D405N;R408S;N440K;V445P;G446S;N460K;S477N;T478K;E484A;F486S;F490S;Q498R;N501Y;Y505H;D614G;H655Y;N679K;P681H;N764K;D796Y;Q954H;N969K;P25-;P26-;A27-;Y144- |

**Supplementary table II:** Neutralizing titers against BA.5, BQ.1.1 and XBB in sera collected from healthy donors in October 2022. For calculations, titers below the minimal dilution of 10 were plotted as 5, and titers at or above the maximal dilution of 640 were plotted as 640.

| **Donors** | **Sex (M/F)** | **Age** | **SARS-CoV-2 infection** | **# of vaccine doses** | **BA.5 $ID_{50}$** | **BQ.1.1 $ID_{50}$** | **XBB $ID_{50}$** |
|---|---|---|---|---|---|---|---|
| 1 | F | 32 | None | 3 | <10 | <10 | <10 |
| 2 | F | 46 | Delta | 2 | 137 | <10 | <10 |
| 3 | F | 44 | None | 3 | <10 | <10 | <10 |
| 4 | F | 30 | BA.5 | 3 | ≥640 | 16 | 16 |
| 5 | M | 38 | None | 3 | <10 | <10 | <10 |
| 6 | F | 56 | BA.5 | 3 | 44 | 14 | <10 |
| 7 | F | 36 | Delta | 2 | 54 | <10 | 10 |
| 8 | F | 26 | None | 3 | 69 | <10 | 10 |
| 9 | F | 29 | BA.5 | 3 | ≥640 | 50 | 29 |
| 10 | F | 54 | BA.5 | 3 | ≥640 | 494 | 113 |
| 11 | M | 45 | BA.1/2 | 3 | 264 | 20 | 14 |
| 12 | F | 28 | BA.5 | 3 | 336 | 151 | 31 |
| 13 | F | 63 | BA.2 | 3 | 165 | 15 | 14 |
| 14 | F | 40 | None | 3 | 180 | 16 | 13 |
| 15 | M | 51 | BA.5 | 3 | 301 | 71 | 23 |
| 16 | F | 58 | None | 3 | <10 | <10 | <10 |
| 17 | F | 36 | BA.1 | 2 | 146 | <10 | <10 |
| 18 | M | 38 | BA.1/2 | 3 | 62 | <10 | <10 |
| 19 | F | 28 | BA.5 | 3 | ≥640 | 86 | 91 |
| 20 | F | 32 | BA.1/2 | 3 | 166 | 12 | <10 |
| 21 | F | 33 | BA.1/2 | 2 | 13 | <10 | <10 |
| 22 | F | 52 | BA.5 | 3 | 621 | 59 | 62 |
| 23 | M | 62 | BA.5 | 3 | 505 | 90 | 22 |
| 24 | F | 54 | None | 3 | <10 | <10 | <10 |
| 25 | F | 35 | BA.1 | 2 | ≥640 | ≥640 | ≥640 |
| 26 | F | 47 | Delta | 3 | 63 | <10 | <10 |
| 27 | F | 50 | None | 3 | <10 | <10 | <10 |
| 28 | F | 51 | BA.5 | 3 | ≥640 | 587 | 369 |
| 29 | M | 56 | BA.1/2 | 3 | 542 | 156 | 14 |
| 30 | F | 28 | BA.1/2 | 3 | 24 | <10 | <10 |
| 31 | F | 46 | BA.1/2 | 3 | 320 | 35 | 59 |
| 32 | M | 39 | None | 4 | ≥640 | 125 | 46 |
